# Recurrent dynamics of prefrontal cortex during context-dependent decision-making

**DOI:** 10.1101/2020.11.27.401539

**Authors:** Zach Cohen, Brian DePasquale, Mikio C. Aoi, Jonathan W. Pillow

**Affiliations:** Department of Computer Science, Princeton University; Princeton Neuroscience Institute, Princeton University; Section of Neurobiology, University of California San Diego; Halıcıoğlu Data Science Institute, University of California San Diego; Department of Psychology, Princeton University

## Abstract

A key problem in systems neuroscience is to understand how neural populations integrate relevant sensory inputs during decision-making. Here, we address this problem by training a structured recurrent neural network to reproduce both psychophysical behavior and neural responses recorded from monkey prefrontal cortex during a context-dependent perceptual decision-making task. Our approach yields a one-to-one mapping of model neurons to recorded neurons, and explicitly incorporates sensory noise governing the animal’s performance as a function of stimulus strength. We then analyze the dynamics of the resulting model in order to understand how the network computes context-dependent decisions. We find that network dynamics preserve both relevant and irrelevant stimulus information, and exhibit a grid of fixed points for different stimulus conditions as opposed to a one-dimensional line attractor. Our work provides new insights into context-dependent decision-making and offers a powerful framework for linking cognitive function with neural activity within an artificial model.

## 1 Introduction

One’s daily experiences are rich with sensory information. Interaction with this volume of data is often guided by a context, an abstract collection of goals, memories and external signifiers that inform appropriate behavioral responses to certain stimuli [1]. Primates in particular possess a remarkable ability to contextually modify behavioral responses to the same stimuli [1–4], demonstrating an ability to flexibly discriminate between relevant and irrelevant information when reasoning about and responding to the external stimuli. Prefrontal cortex (PFC)—a brain area implicated in executive functioning [1], executive control [5], and a wide range of abstract, high-level reasoning tasks [3, 6–8]—is thought to play a central role in context-dependent perceptual reasoning [1–3,8, 9]. The exact mechanism underlying PFC’s contribution to such reasoning, however, remains an open question [4, 10–12].

Artificial cognitive neural network models allow researchers to generate testable hypotheses concerning the functional neural mechanisms underlying complex behaviors [4, 13–18]. Recent work has employed artificial network models to try and understand the neural mechanisms underlying context-dependent perceptual reasoning [4, 13]. This work is largely task oriented: The dynamics of neurons from a network trained to simulate a task of interest are studied in order to gain insight into how the brain may solve the same task [4, 13]. Task-based methodology affords a hypothesis of how the brain performs some computation [4, 13, 14, 19], but leaves open a critical question of whether or not the trained artificial neurons accomplish the task of interest using the same—or even similar—computational techniques as those employed by their supposed biological analogues.

Here, we considered two hypotheses. The first asks if it is feasible to constrain the internal dynamics of a computational model to approximate those recorded in neurological data in addition to training the model to solve a context-dependent perceptual discrimination task. To test this hypothesis, we trained a recurrent neural network (RNN) to solve a context-dependent perceptual discrimination task while simultaneously training the internal dynamics of the network to simulate those observed in experimental recordings of rhesus monkeys trained on the same task. We found that it is possible to train artificial models to achieve both near-perfect and more biologically realistic performance on perceptual discrimination tasks when the internal dynamics of the network are set such that there is a one-to-one correspondence between network neurons and neurons from experimental recordings.

The second hypothesis asks if models with dynamics constrained to fit biological data compute a perceptual choice in a manner that is similar to that employed by networks trained without such constraints. To test the second hypothesis, we “reverse-engineered” [20] the network model to understand how it simulates observed neural dynamics and how these dynamics sub-serve context-sensitive behavior. We then compared our results to those obtained using similar methodology on a task-based network. We identified a notable difference between the computational mechanism employed by our model and that employed by task-based models in solving the perceptual discrimination task. Previous task-based models of flexible perceptual discrimination have hypothesized that irrelevant information is dynamically suppressed in neocortical circuits in converging upon a perceptual decision, and that relevant information is integrated along a line attractor [4]. In our model, by contrast, both relevant and irrelevant information are persistently represented and integrated across a planar manifold in the neural state space throughout the course of making a perceptual decision. This suggests that a more complex computation than previously hypothesized underlies flexible perceptual reasoning.

Our results challenge past mechanistic models of context-dependent perceptual decision-making. Still, our results prove to be more consistent with more recent work aimed at understanding the computational mechanism employed by PFC in performing context-dependent perceptual reasoning [21]. Our findings suggest that the addition of dynamical constraints helped our model identify a biologically realistic solution to context-dependent perceptual decision-making.

## 2 Results

The model that we use is a randomly initialized neural network that is comprised of two distinct populations, which we term “hidden” and “observed” populations (described in detail in Appendix C). Observed neurons are recurrently connected and are trained (Appendix C.2) to simulate experimental neurological recordings associated with some contextually coded set of stimuli. These stimuli are introduced to the network via the hidden population. The hidden population connects to the observed population by unmodified feed-forward connections. Inputs are applied to the hidden population. The observed population receives input by internal recurrent connections and from the input population. The hidden population amplifies and low-pass filters the inputs, and then conveys them to the recurrent population. The hidden population can be thought of as a simplified abstraction of upstream brain areas, like primary visual cortex, through which external information is first encoded into neural signals [22] before reaching downstream cortices, like PFC (which the observed population is designed to model).

Critically, we constrain the dynamics of the observed population by training it to simulate experimental recordings of FEF of rhesus monkeys taken while these monkeys performed a perceptual discrimination task (schematized in *Figure 1*.a,b, data from Mante et al. [4]). The readout of any neuron in the observed population is its firing rate at any time *t* (*Figure 1*.c), and this read-out is trained to continuously match the instantaneous firing rate of its corresponding “neuron” from experimental recordings. In other words, each observed neuron is mapped one-to-one to a single experimental neuron (Appendix C).

**Figure 1:**
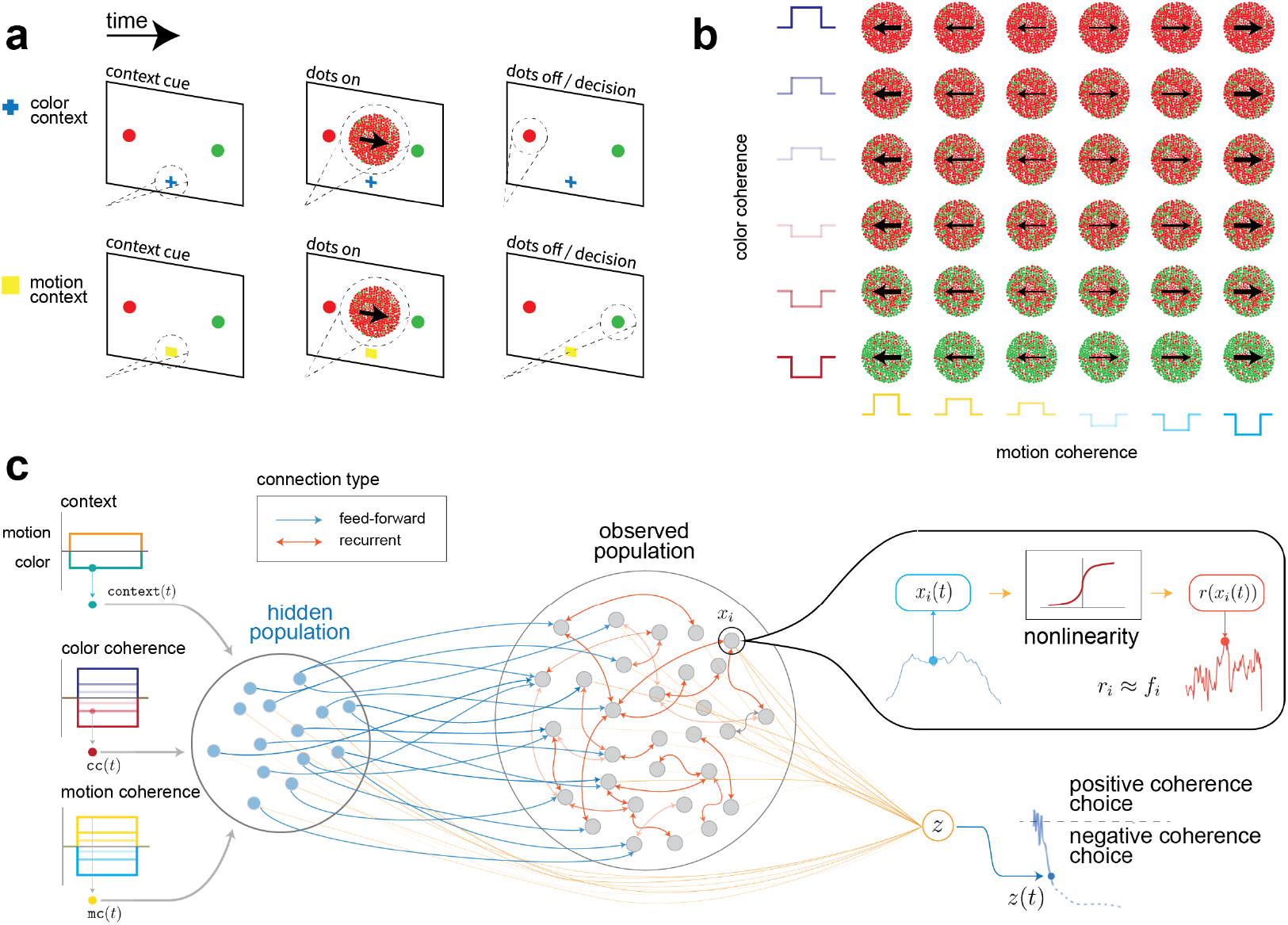
Perceptual discrimination task and model schematic. **a.** A time course of the dot-discrimination task, with each phase labeled. In the original task, the monkeys were presented with a rendering of randomized dots on a display, which, in each trial, had some motion (drifting to the left or right) and color (primarily red or green) coherence. In each trial, the monkeys were presented with a context cue, instructing them to focus on and later report with a saccade either the color or motion coherence of the dots. During this task, neural recordings were taken from the FEF. These recordings were taken at the population level of neurons in the FEF. **b.** A set of possible color and motion coherence combinations of the dots on any trial along with the corresponding task variables used by our model. **c.** A schematic of our model. The network is split into observed neurons, whose dynamics are constrained by experimental data, and hidden neurons. The observed state of the network is the firing rate of each neuron in the observed population. A behavioral node (*z* in the schematic) integrates the firing rates of each neuron and reports a behavioral task variable (either reporting +1 or −1 depending on the coherence values of the relevant input stimuli).

We modeled the task variables by coding all possible combinations of stimulus coherence values used in the discrimination task with a binary representation of a “context”, indicating to which sensory stream the network should attend. A final readout neuron indicates the behavioral choice made by the network. The behavioral node’s readout is the weighted sum of the instantaneous firing rates of the neurons in the network at each time step. When the coherence value of the relevant stimulus (as indicated by the context cue) is positive, the behavioral node is trained to tend towards a positive value. The converse is true when the relevant stimulus takes on a negative value.

### 2.1 Recurrent network model dynamics can be constrained to fit contextual neural activity

We verified that the network’s observed population accurately captures experimentally recorded neural dynamics on a per-condition basis (i.e., the network’s observed neurons exhibit the same trajectories as experimentally recorded neurons when the monkey and the network are presented with the same stimuli).

After training, observed neurons in our model follow trajectories that qualitatively resemble target experimental recordings, even when strong noise is injected onto the input stimulus to which these experimental recordings are associated (*Figure 2*.a and *Figure 2*.b, top rows of each panel). Across the strengths of the irrelevant stimulus, neural trajectories in a context are separable according to the corresponding choice. The network captures this pattern, while demonstrating the ability to capture the relatively small but crucial differences between neural trajectories within the same choice but driven by different relevant stimulus strengths.

**Figure 2:**
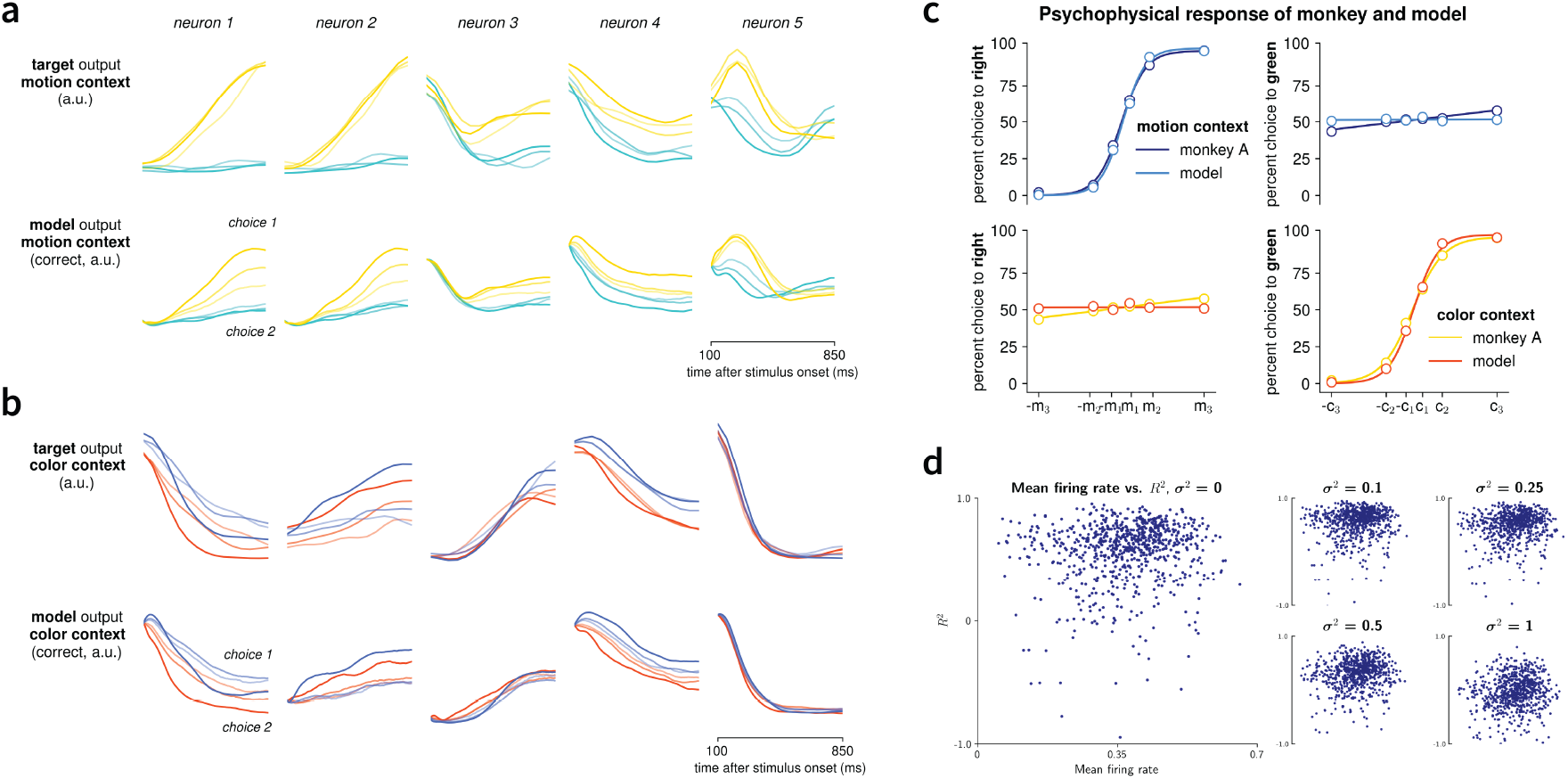
Result of simulating experimental data using our model. **a.** The top row shows the target PSTH, the second row shows the model PSTH on trials in which the network correctly reported the direction of the relevant stimulus. Each subgraph on a row depicts a single neuron’s response, averaged over all irrelevant color stimulus values for each motion coherence value. Gold indicates positive motion coherence, cyan indicates negative coherence; opacity is proportional to the coherence strength. **b.** The same as **a.** but in the color context. Blue indicates “positive” color coherence, red indicates “negative” color coherence. Annotations on the trace indicate the associated behavioral choice of the stimulus driving a particular neural response; *choice 1* indicates a positive coherence response, and *choice 2* indicates a negative coherence response. In experiments here, motion and color signals were corrupted by Gaussian noise drawn i.i.d from 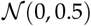. Responses are averaged over 150 trials. Neural responses of the network when only presented with the context cue that precede these traces are not shown. **c.** Psychophysical performance of the network, presented alongside the psychophysical performance of monkey A from [4] to demonstrate the similarity in behavior. The top panel presents discrimination in the motion context, and the bottom panel demonstrates discrimination in the color context. Performances was measured over 17, 280 trials, and the results here are the result of inference decisions made by the network when the stimulus signal was corrupted with Gaussian noise with variance *σ*^2^ = 0.5. Increasing the stochastic noise that corrupts the stimulus signal negatively impacts the 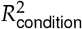 and 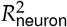 statistics, as expected. Noise more strongly degrades performance of neurons in the network simulating experimental neurons whose mean firing rate is less than 0.5, while neurons with mean firing rates greater than 0.7 remain consistent under higher degrees of noise corruption. At the highest levels of noise, some neurons in the population demonstrate complete anti-correlation with their respective target functions. Still, the average 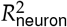 across all neurons decreases modestly across different noise values. **d.** The mean firing rate of observed population neurons is plotted against 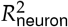. As the variance of the Gaussian noise corrupting the input stimulus. *σ*^2^, increases, 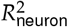 across neurons decreases.

We use the 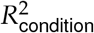 and 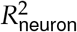 statistics (Appendix D) to quantify the network’s accuracy in simulating recorded dynamics. We relate this metric to the mean firing rate of a neuron over the course of a trial. *Figure 2*.d shows several plots that relate 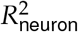 to noise injected on the stimulus and the mean firing rate of neurons in the model. There exists a weak positive correlation (*r*^2^ = 0.1437, *P* = 3.55 × 10^−5^) between neuron firing rate and its 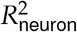 for simulations that do not involve stimulus corruption.

### 2.2 Models constrained with experimental data exhibit biologically realistic behavior

Our model links the psychophysical response in the experimental regime with the neural codes that underpin it. *Figure 2*.c shows that the model is able to achieve the psychophysical performance of the monkey from which the neural and behavior data is taken, matching the behavioral curve that demonstrates preferential selectivity for the relevant stimulus in some respective contextual regime (top-left and bottom-right panels) as well as indifference to markers indicating response variables from the irrelevant sensory stream (the bottom-left panel and the top-right panel).

The psychophysical response demonstrates a key aspect of a realistic form of stimulus integration: When the relevant stimulus has a high coherence value, we expect an integrator to be able to reliably report salient information about the signal, even in the presence of noise. As the signal decreases in coherence, we expect the integrator to make more mistakes; as the mean coherence reaches zero, particularly when noise is applied to the signal, it becomes more difficult to accurately report the sign of the mean coherence value.

The psychophysical behavior demonstrated in *Figure 2*.c is induced only in trials in which the stimulus is corrupted by noise. In trials with no stimulus corruption, the network is able to perfectly discriminate sensory evidence, even at low coherence values (Appendix B.1). The behavior of the network in this case is largely consistent with previous results that examine network simulation of only the behavioral response [4].

Particularly in networks with a large number of hidden neurons, high noise corruption shaped the psychophysical response of the network to generate biologically realistic feature selection without severely impairing the ability of the observed population to reliably simulate experimental recordings. Previous iterations of our work either did not employ hidden filtering neurons, or treated hidden and observed neurons entirely the same (Appendix A). Both of these prior approaches failed to simultaneously simulate the target biological characteristics in both the neurological and behavioral modalities. Of course, simulating experimental psychophysical behavior at the expense of the ability to simulate observed biological neural dynamics defeats the purpose that opens the explorations in this work in the first place. Conversely, simulating just neurological dynamics at the expense of associated target behavior tells us only about the dynamics of neural circuits alone, and less how those dynamics give rise to behaviors of interest.

### 2.3 Hidden population dynamics

Much of the accuracy achieved by the model in representing both behavioral and neuronal targets can be attributed to the model structure; in particular, the way in which information is input into the network via the hidden population (see Appendix C and *Figure 3*). Beyond helping the network in accomplishing the context-dependent perceptual integration task and in filtering out noise, additional hidden units increase 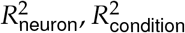 across simulations, and also increase the model’s resistance to noise corruption, both applied directly to the neurons (see *Equation 3.1*) and to the color and motion stimuli, as demonstrated in *Figure 3*.d.

**Figure 3:**
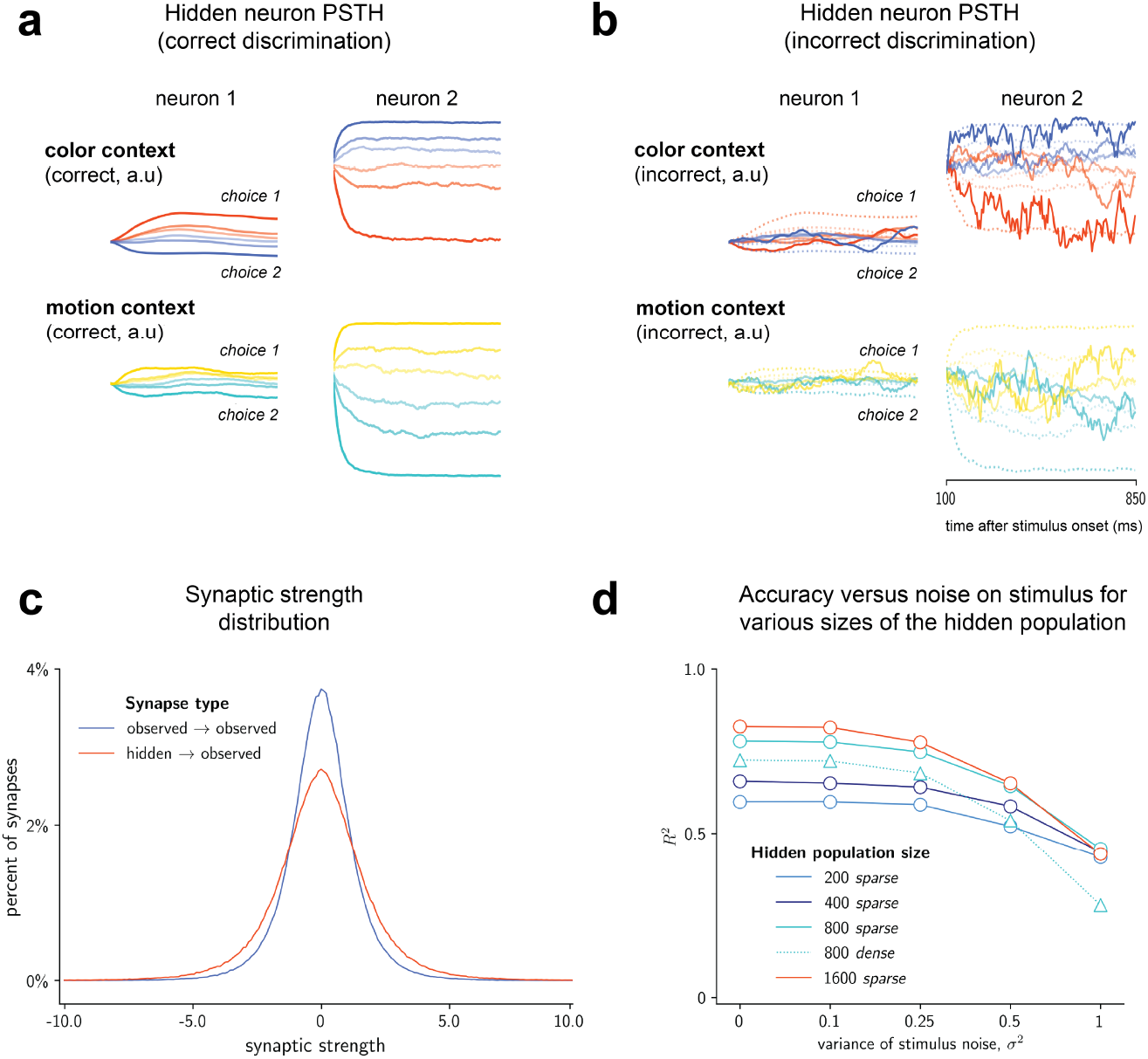
**a.** PSTHs of sampled hidden neurons from the model population. The top row displays responses in the color context, and the bottom row displays responses in the color context. The color coding used here is the same as that used in *Figure 2*. **b.** PSTHs from sample neurons in the hidden population recorded on trials in which the network reported the incorrect value of the relevant stimulus coherence. The solid lines indicate the response PSTH on trials in which the network reported the incorrect coherence value, and the dotted lines indicate the response PSTH of the same neuron on trials of correct discrimination in the color context (top row) and motion context (bottom row). The color scheme used here is the same of that used in *Figure 2*. **c.** The synaptic weight distribution between neurons exclusively in the hidden population (blue), and between neurons projecting from the hidden population to the observed population (red). **d**. The strength of stimulus noise corruption—quantified by the variance of the Gaussian distribution from which the noise was sampled, *σ*^2^—against the median 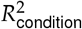 across all conditions for networks with different numbers of hidden neurons in the population. The dotted line with triangles represents these measurements with one of the networks not used in this study: the alternative network model in which the input stimulus is projected onto the whole population (we term this network “dense” because of the density property of the input matrix used by the network).

The dynamics of the hidden neurons follow a surprisingly simple schematic. A significant portion of neurons in the hidden population saturate at the tails of the sigmoid nonlinearity (*r*(·)), tending towards and staying at either 0 or 1 for the majority of the trial. The ones that do not saturate appear to deflect from the mean firing rate of the neuron across trials at a magnitude proportional to the strength of the input stimulus over the first 10-20 ms of the trial. For the remainder of the trial, the firing rates of these neurons remain steady, as demonstrated in *Figure 3*. This behavior of the hidden neurons would suggest the existence of several attractors towards which the hidden population quickly travels within the state space of the network. The long-term behavior of the network in each trial suggests that these attractors are stable, and that the gradients around these attractors are quite steep.

The neurons in the observed population do not receive direct projections of the stimuli; in fact, they are only “aware” of fluctuations within the dynamics of other observed neurons and hidden neurons. This would suggest that the hidden neuron population’s contribution to solving the task involves steadily representing the input stimulus over the course of the trial. Replicating the exact value of the stimulus is not necessary, as there need only exist a relative difference between all coherence value representations in order for the network to be able to complete the task (recurrent weights and weights presynaptic to the readout node can have positive or negative sign of varying magnitude).

*Figure 3*.c shows a histogram of the magnitude of synaptic weights between neurons in the observed population and between neurons in the observed and hidden populations^1^. The mean strengths of these two subsets of synaptic weights are both 0, but the variance is much wider over the weights projecting from the hidden population to the observed population. In other words, any given neuron in the observed population receives a stronger driving force from the hidden population than from other neurons in the observed population. Put differently, one can think of each observed neuron as, at least on an individual basis, preferentially strengthening its connection with any given hidden neuron, on average, as opposed to any given observed neuron. Persistent representation of a given stimulus in the observed population, then, relies at least in part on the representation of that stimulus in the hidden population.

The PSTHs of the hidden neurons on trials in which the network reported the incorrect choice variable also help in characterizing the role of hidden neurons in representing task variables. *Figure 3*.a depicts PSTHs from the first two neurons in *Figure 3*.b on failed trials. On failed trials, the magnitude of the deflection of the PSTH away from the mean firing rate of a neuron at the start of the trial is nowhere near the strength of the corresponding deflection on trials in which the network correctly solved the behavioral portion of the task. Beyond that, the PSTH for any given stimulus strength tends to drift back and forth in firing rate space, concentrating towards the end of the trial somewhere near the mean firing rate of the neuron across different stimulus strengths in correct trials. This would suggest that incorrect discrimination is a result of the hidden population being unable to represent stimulus information correctly insofar as the network is trained to expect over the course of a trial.

These observations collectively point towards a relatively simple characterization of how hidden neurons help promote correct simulation of target PSTHs and associated behavioral variables. The task of correctly representing contextual information involves learning recurrent connections within the network that give rise to context- and stimulus coherence-specific attractors within the state space spanned by the network. Stimulus inputs drive the hidden population towards the respective attractors learned during training that correspond to the coherence value and sign of the relevant *and* irrelevant stimuli.

### 2.4 Tracing stimulus-driven network trajectories in lower dimensions

In order to understand *how* our model solves the context-dependent perceptual discrimination task when its internal dynamics are trained to simulate experimental data, we turned to “reverse-engineering” [20] our network. This methodology (Appendix C.5) involves identifying fixed and slow points in the phase space spanned by the optimized network, and characterizing—both quantitatively and qualitatively—local dynamics in the neighborhoods of these fixed points. Features of local attractor dynamics of the network, from the spatial organization of fixed points to the stereotyped trajectories of the network around stable points in response to certain inputs, can offer global insight into how the network solves tasks of interest.

We found two different sets of fixed points in our model’s state space. One set was found by identifying regions of stable dynamics while driving the network with only the contextual cue as input, and the other set was found by driving the network with the contextual cue as well as one of the 36 possible combinations of color and motion stimuli as input on any given run of the optimization routine (*Figure 4*.a). The spectral qualities of the Jacobian of the network at a given fixed point can offer a quantitative perspective of network activity when in the fixed point region [4, 16, 23, 24], and so we performed an eigendecomposition of the Jacobian of our network at the fixed points in its phase space (*Figure 4*.b left panel).

**Figure 4:**
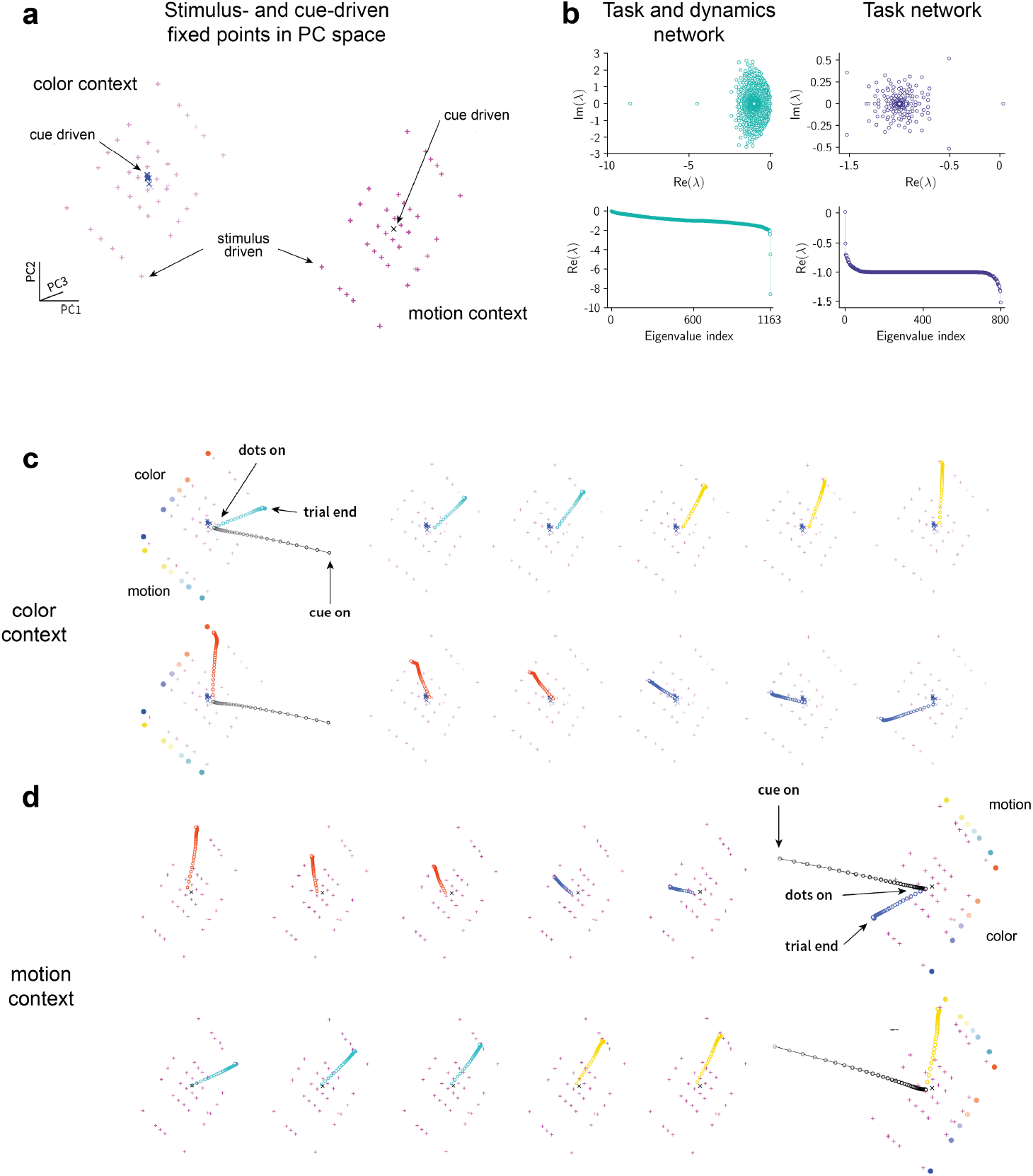
**a.** Stimulus- and cue-driven fixed points in the state space of the network. We use the top three principal components from sampled activations of the network here to visualize the state space in 3 dimensions. Magenta crosses are stimulus-driven fixed points in the motion context and pink crosses are stimulus-driven fixed points in the color context. Clustered points at the center of these planes are the respective context-cue driven points. **b.** Eigenspectra of fixed points in our model network (left side, top and bottom) and of another model trained to perform just the perceptual discrimination task (right side, top and bottom). See Appendix C.3 for details on the structure and training method used for this model. See *Figure 9* in extended figures for representations of the population activity of the network trained to perform just the perceptual reasoning task within the state space as defined in [4]. **c.** Network trajectories in the color context. For the trajectories in the top row, the color coherence of the stimulus presented to the network is held constant, and the coherence value of the motion stimulus is varied over all 6 possible values and for the trajectories in the bottom row, the motion coherence value is held constant while the coherence value of the color stimulus is varied over all 6 possible values. **d.** The same as **c**, but in the motion context.

The eigenspectra of the Jacobian of the network at the fixed points we identified all followed a common schematic (*Figure 4*.b, left panel). The largest real part of the eigenvalues is at or close to 0. The remaining eigenvalues descend in magnitude fairly slowly—the real parts of 86% of the eigenvalues, on average, are greater than −1. In general, between two and five eigenvalues have significantly larger real parts, typically between −4 and −12. This suggests that in regions of stability in the state space, local perturbations of the network induced by external inputs evolve slowly over many hundreds of dimensions. In other words, information is integrated and represented by the network across hundreds of dimensions of the state space—the distributed integration of this information is a critical component of the network’s ability to both simulate experimental neural dynamics in addition to simulating the expected behavioral sensitivities to context when discriminating relevant and irrelevant stimuli.

The fixed point eigenspectra we identified are fairly different than those of task-based network models. Networks trained to solve just the perceptual discrimination task by reporting the sign of the relevant stimulus, for example, reveal a common spectral organization [4] (see *Figure 4*.b, right panel for the eigenspectrum of a network we trained to solve just the discrimination task, and Appendix C.3 for more information about that network). The eigenspectrum of the Jacobian of the network at any of the fixed points in the state space has a unitary eigenvalue, which corresponds to the dimension along which inputs are both projected and persist within the network’s phase space [4]. The remaining eigenvalues have fairly large negative real parts, and inputs projected along the dimensions corresponding to these non-unitary eigenvalues quickly decay out of the system.

The discrepancy in the eigenspectra of fixed points in our network and in task-based networks suggest that these two networks are accomplishing the same context-dependent perceptual discrimination task differently. Whereas in task-based networks, input integration happens along a drastically simplified subspace (a line attractor, for example) of the larger network state space, our model represents information in a significantly higher-dimensional fashion over the course of a trial.

Previous work using task-based networks took advantage of the lower-dimensional nature of the network’s solution to the task, projecting network activity in response to experimental stimuli into a subspace spanned, in part, by the dominant axis associated with the expected unitary eigenvalue [4]. Observing the network’s dynamics in this reduced (and human-visible) sub-space offers intuition for how the network operates in a higher-dimensional sense.

In lieu of estimating a projection subspace using the unitary integration axis, we projected the fixed points we found into the subspace spanned by the top 3 principal components over a collection of recorded activations. This method of lower-dimensional analysis is common in the reverse-engineering literature [16, 18, 19]. In our case, 99% of the variance in network trajectories is accounted for in the top three principal components. We found that the fixed points are organized in a very specific structure (*Figure 4* .a). The fixed points found by driving the network with the stimulus clamped on are organized in what appear to be two plane attractors that are separable by context. Each contextual plane embeds 36 fixed points, one for each possible combination of color and motion coherence values. The space between each fixed point in principal component space is proportional to the magnitude of both the relevant and irrelevant stimulus.

Fixed points that were found by driving the network with just a context cue are also separable by context within the state space. They are the center of the planes along which the stimulus-driven fixed points lie for their respective contexts.

While examining the fixed points within the state space spanned by the network provides intuition for how the network solves the task of context-dependent feature selection, this tells only a small portion of the story. We proceed to further characterize the computations performed by our network model by projecting recorded activations of network neurons from simulations across all stimulus conditions. We map how the network represents its state, and thus, the information it has integrated, via its location in the state space.

A trial of context-dependent perceptual inference follows a fairly simple schematic, which is illustrated in *Figure 4*.c-d. When the network is presented with a context cue informing it of which oncoming stream it should attend to, the network state quickly drifts in the state space towards the set of fixed cue-fixed points embedded within the attractor plane corresponding to the relevant feature (determined by the context cue). The network stays in this area until the “dots” (the sensory information corresponding to color and motion) come on, and the network state quickly moves in the direction of the stimulus-driven fixed point on the attractor plane corresponding to the combination of coherence values of both the relevant and irrelevant sensory stream presented on that trial. Over the course of the trial (as long as the input is clamped on), the network stays tightly bound to the stimulus-driven fixed point. When the stimulus turns off, the network returns to resting state, drifting back in the state space to the point at which it began the trial.

The trajectories of the network in this subspace, as well as the existence of fixed points on either context plane corresponding to all permutations of relevant and irrelevant cues, are telling. The behavior of the network, specifically its coordinated projection that is sensitive to *both* the relevant and irrelevant data for a given trial, suggests that no information, neither relevant or irrelevant, is dynamically deleted during the process of input integration. In both contexts, the network represents input information along something like a plane manifold (although, we could easily imagine that if there were, say, three inputs to integrate, integration would take place along a higher-dimensional manifold) with orthogonal planar axes representing color and motion coherence values. The context-sensitive response of the network is a result of a highly separable state space according to context, with trajectories in either context represented by location along a contextual attractor manifold. Because these manifolds are separable along a context, location on either manifold can represent different context-dependent neural trajectories, while not requiring that *only* relevant sensory data drive the network to some position on the manifold.

## 3 Discussion / Conclusion

In this work, we show that an artificial recurrent neural network can be trained to solve a context-dependent perceptual discrimination task when its internal dynamics are constrained to fit electrophysiological data recorded from live subjects performing the same type of feature discrimination.

Other work has employed artificial models to simulate neural data [25, 26]. A distinguishing feature of our model is that the readout of any neuron is simply the firing rate of that neuron at any time, rather than a weighted summation of the firing rates of neurons presynaptic to it [25]. This is a subtle but important distinction: it means that, at least in the observed population, the dynamics of each neuron in closely mimic those of experimental data. A deeper consequence of this is that our model offers direct insight into how each neuron in the population contributes to the computations that constitute contextual sensitivity to stimuli.

In contrast to work from [25, 26], we saw that simple recurrent connections within the observed population are insufficient in giving rise to the observed neural firing patterns from data, and that a hidden population whose dynamics are not fixed to a target function are necessary in helping the network generate the appropriate target dynamics. This is likely a result of enforcing a direct correspondence between neurons in the artificial population and those in the recorded neural circuits.

Other work has trained networks to solve tasks while also training its internal dynamics during training [27]. This direction is fairly new, and, at least in the example cited here, relies on simulated neurological data as a target for the trained dynamics of artificial models. In contrast, the model we present here uses live recorded neurological data as the target set of dynamics of observed artificial neurons, drawing a direct computational correspondence between experimental neurological data and observed behaviors associated with that data.

Simulation of neural activities recorded in experimental settings provides insight into how the codes themselves are generated, but it leaves unresolved the connection between neural activity and the associated observed behavioral activity. We resolve that link by incorporating a behavioral node within our model whose dynamics are similar to that of an ordinary neuron in the population. This node integrates the information simulated by the observed and hidden populations to generate a target behavioral choice. The node responsible for reporting the coherence of the relevant sensory stream makes a choice regarding the stimulus using only information as the network represents it.

The computational mechanism governing our optimized network complicates the schemes proposed in related work that employs task-based computational models [4, 13, 19]. The spectral distributions of the network around fixed points, unlike the corresponding distributions of task-based networks, suggest that when artificial neurons are held to simulate experimental ones, integration of input information in the network state space necessarily evolves along hundreds of dimensions. Investigating a lower-dimensional subspace spanned by the axes along which maximal variance of the network response is captured shows that while the integration manifold may span several hundred dimensions, it can be approximated in only two dimensions spanned by axes corresponding to the relative coherence strengths of both motion and color stimuli. Movement along this plane during integration of sensory evidence demonstrates that the system retains information about both the relevant and irrelevant streams, rather than dynamically deleting irrelevant data from the state space.

It is perhaps not surprising that the integration of stimuli in our model follows a higher-dimensional, more complex scheme. The target function for the observed population, while highly correlated between neurons in the population, maintains distinct target functions for all permutations of relevant and irrelevant sensory data. Therefore, it is absolutely necessary for the network to find a way to represent both relevant and irrelevant sensory data, even if the corresponding choice is not driven by the irrelevant data.

The very existence of distinct PSTHs for varying coherence values of an irrelevant stimulus when the relevant stimulus coherence value is held fixed is some indication that irrelevant sensory information *is* in fact encoded within the hypothetical state space in which actual neurons compute information. Recent work that proposes alternative schemes for reducing the dimensionality of the same dataset used by Mante et al. [4] has effectively demonstrated that irrelevant sensory data is certainly encoded in recorded neural activity [21]. In fact, this work demonstrates that both the relevant *and* irrelevant sensory data can be decoded from the PSTH for trials of both correct and incorrect discrimination [21]. These findings are far more consistent the activity of our network than the model from [4], as, in making any behavioral choice, ours must represent both the relevant and irrelevant sensory data in the state space.

One possible critique of the set-up and conclusions drawn here rests on the number of degrees of freedom in constructing artificial models, from choosing a non-linearity to tuning the sparsity of recurrent connections between neurons [19]. Recent work, however, has demonstrated at least partial universality in the ways in which RNNs learn to simulate complex dynamics in spite of respective differences in training methods and other internal features [19]. Thus, we might conclude that the mechanistic aspects uncovered by our “reverse-engineering”—*how* the network implements a solution to the task—may generalize beyond the variety of possible parameters we can tune, and hints at a deeper computational mechanism shared by circuits of neurons in the brain and ensembles of artificial neurons trained to the dynamics of these circuits.

In increasing the complexity of the task performed by the model network, and in increasing the biological realism of the model (up to an asymptotic degree imposed by the very type of model we used), we saw that the low-dimensional characterization of the functional mechanism underlying context-driven feature integration and selection also grew in complexity. Still, through use of techniques similar to those employed in studies collectively used by this investigation as a starting point, we were able to surmise a coherent, semi-quantitative description of a possible mechanism underlying context-dependent computation in artificial circuits.

## A Alternative network structures

**Figure 5:**
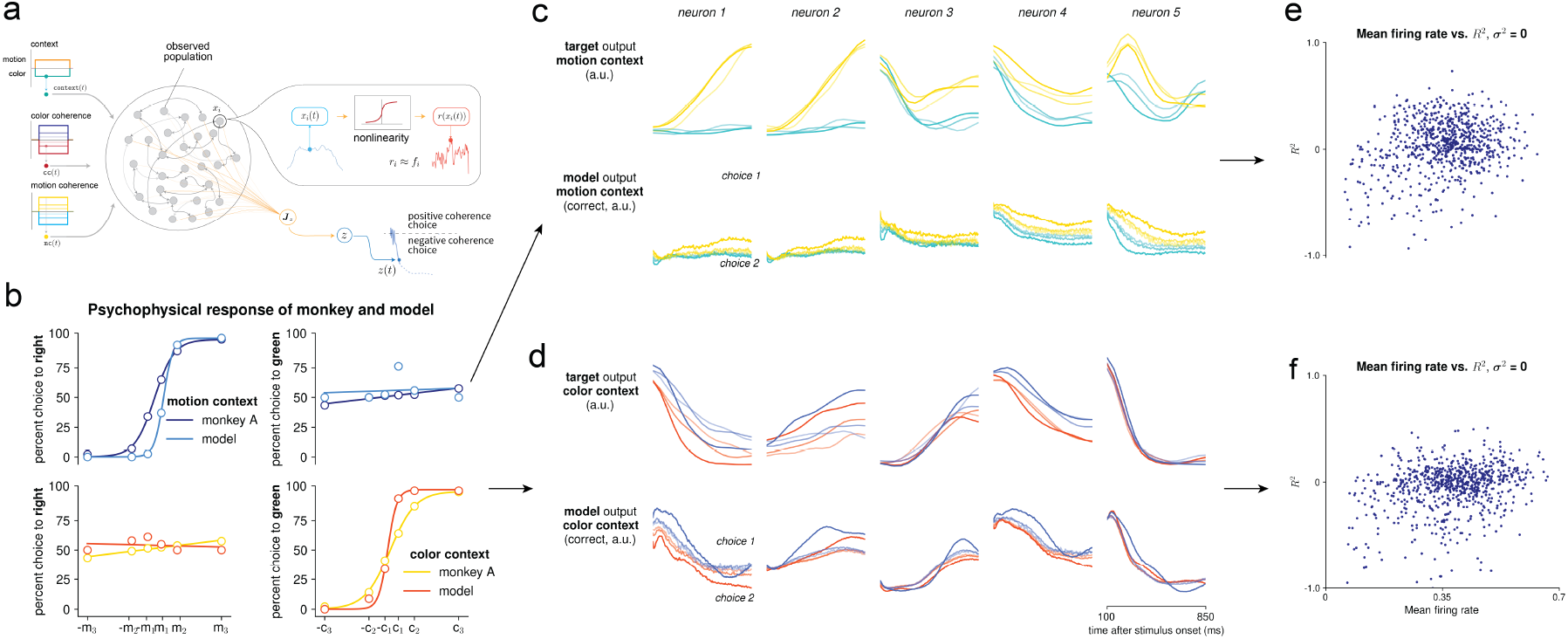
**a.** A network model without hidden neurons. **b.** is the psychophysical performance of the network alternative in **a.** when the evidence stimulus is corrupted by noise drawn i.i.d from 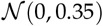. The top row of **c.** is the target output of neurons 1-5 in the motion context, and the bottom row is the output of the network when the evidence stimulus is corrupted by noise drawn i.i.d from 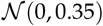. **d.** is the same as **c.**, but in the color context. **e.** and **f.** depict the relationship between the mean firing rate of neurons sampled from the network population and an individual neuron’s 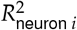 in the motion and color contexts, respectively.

Our model design treats hidden and observed neurons quite differently. Not only do observed neurons have a target function, but the structure of the network is largely centered around a meaningful differentiation between the two sets of neurons: Input stimuli are only projected onto the hidden neurons, and synapses presynaptic to the hidden population are never modified, while all synapses (both from observed and hidden neurons) presynaptic to the observed population are modified during training. This model design is largely a result of several explorations of other model designs which achieved significantly worse performance than the model that is the main focus of this work. In justifying our model structure, we compare our model of choice and two others built and surveyed in the course of this work. One has no hidden neurons in the population (*Figure 5*.a), and in another, inputs are projected onto the entire population (*Figure 6*.a). These comparisons also point to crucial ingredients in our model for generating flexible neural dynamics.

*Figures 5* and *6* show both the model structures of the two alternatives, as well as two sample PSTHs recorded from simulating the respective models. The model without any hidden neurons demonstrates a poor ability to capture the the dynamics of the neurons in the population beyond the discriminability of the PSTHs of the recorded population at the level of behavioral choice. The psychophysical performance of this model is also not on par with that of the model that is the focus of this paper, with behavioral choices showing limited correlation with the strength of a stimulus in its respective contextual regime (*Figure 5*.b). The inputs to the behavioral node are simply the neurons presynaptic to it in the network, and therefore its dynamics are solely driven by the dynamics of the observed neurons. The poor psychophysical performance of this model, then, appears to be a natural extension of the model’s poor performance in simulating neural trajectories.

**Figure 6:**
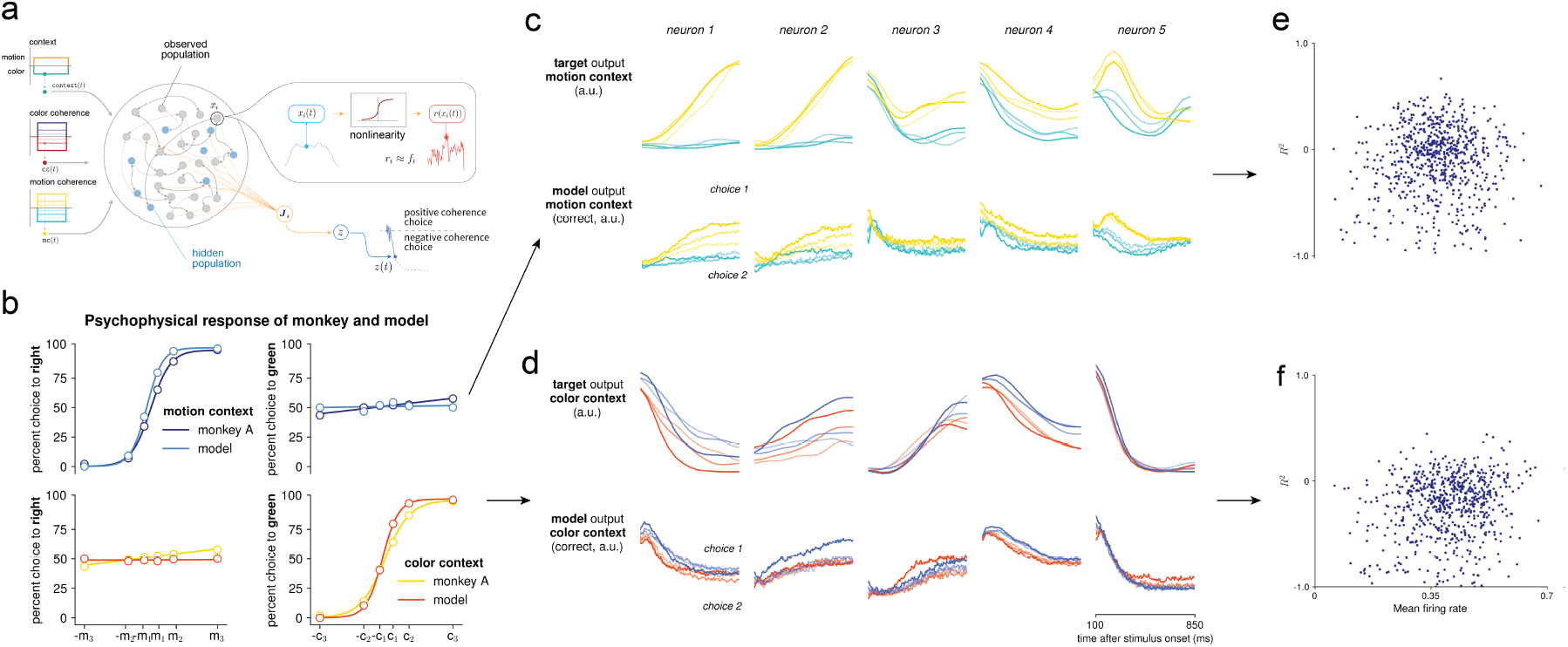
**a.** A network model in which stimuli and context cues are projected to the entire population, and hidden neurons are recurrently connected both to themselves and to the observed population. **b.** The psychophysical performance of the network alternative in **a.** when the evidence stimulus is corrupted by noise drawn i.i.d from 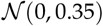. The top row of **c.** is the target output of neurons 1-5 in the motion context, and the bottom row is the output of the network when the evidence stimulus is corrupted by noise drawn i.i.d from 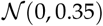. **d.** is the same as **c.**, but in the color context. **e.** and **f.** depict the relationship between the mean firing rate of neurons sampled from the network population and an individual neuron’s 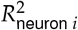 in the motion and color contexts, respectively.

At high levels (*σ*^2^ ≥ 0.25) of stimulus corruption with stochastic noise, distinction of the neural trajectories based on choice completely degrades in the model with no hidden neurons. Unlike the model with no hidden neurons, a model with hidden neurons, but in which the input stimulus is equally projected onto the whole population, shows a strong ability to capture neural dynamics across conditions beyond just discrimination at the level of choice. This would suggest that despite the fact that hidden neurons are not designed to approximate a target function, optimizing the recurrent weights of the network relies on the long-term steady state of hidden neurons; weights presynaptic to the observed population from the hidden population are adjusted accordingly, and the hidden population largely facilitates the observed population in simulating target trajectories. Further, this suggests simple recurrent connectivity between only the observed neurons from the experimental data cannot give rise to the dynamics observed in the recordings that comprise the target dataset in this work. Likely, other circuits of neurons in the brain are critical in producing the trajectories observed in experimental data, and also in generating the associated target behavior.

While the model with hidden neurons mixed in the population is capable of capturing neural dynamics on trials in which the stimulus is not corrupted by noise, injecting strong stochastic noise into the stimulus dramatically impairs performance. We might be satisfied with this type of result, as it seems like the logical consequence of our methodology. These results, however, pose two issues: First, variance on the noise of an incoming sensory signal into the brain have little to no effect on the amount of noise in the PSTHs of cortical neurons. In attempting to model neural circuits with as much realism as possible, the behavior of the model when presented with noisy stimuli would be a major barrier. The psychophysical response of the network is perfect— and thus, not biologically realistic—in experiments with no signal corruption, but more closely approximates experimental psychophysical response of the monkey. This points to the second issue: In this model, there exists only a tenuous link between PSTHs of the observed neurons and the behavioral output of the network. We suppose by design that the trajectories of the population largely (if not entirely) motivate the selection of the appropriate behavioral variable. In particular, we suppose that the neural recordings from our target dataset *specifically* are the neural representations of test stimuli and that they give rise directly to the psychophysical response of the monkey. A model that matches psychophysical response, but whose observed trajectories bear little likeness to the target trajectories, would not be particularly useful in determining the underlying nature of the link between neurological and psychophysical activity in context-dependent tasks.

The primary model we use in this work is free of these issues. In fact, even in the trials with no stimulus corruption, it achieves a higher 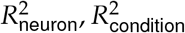 than the either of the other models. This is likely because observed neurons in both of the models not used in this work must integrate both the direct projection of the stimulus in addition to the firing rates of all other neurons in the network at each time step.

## B Additional Figures

## B.1 Network models are perfect integrators without noise on the signal

**Figure 7:**
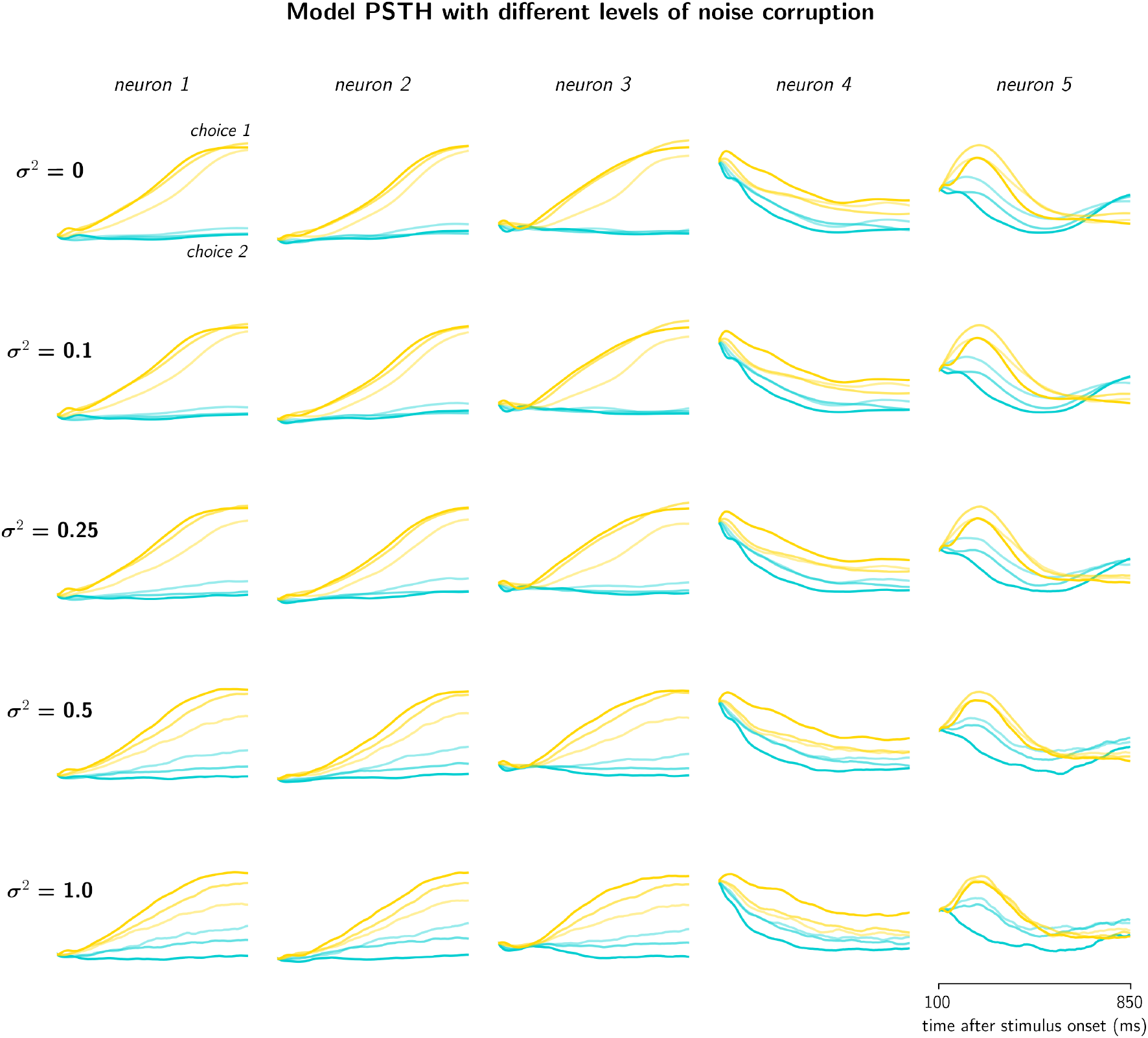
Model PSTHs with different levels of noise corruption. . Recorded PSTHs in this figure are taken in the motion context, with the same color scheme as that in *Figure 2*. Noise corruption does impair general simulation performance, but not nearly to the degree that it does in other candidate models, as seen in *Figure 5*.

## B.2 Psychophysical response of candidate models with and without noise corruption of the stimulus

**Figure 8:**
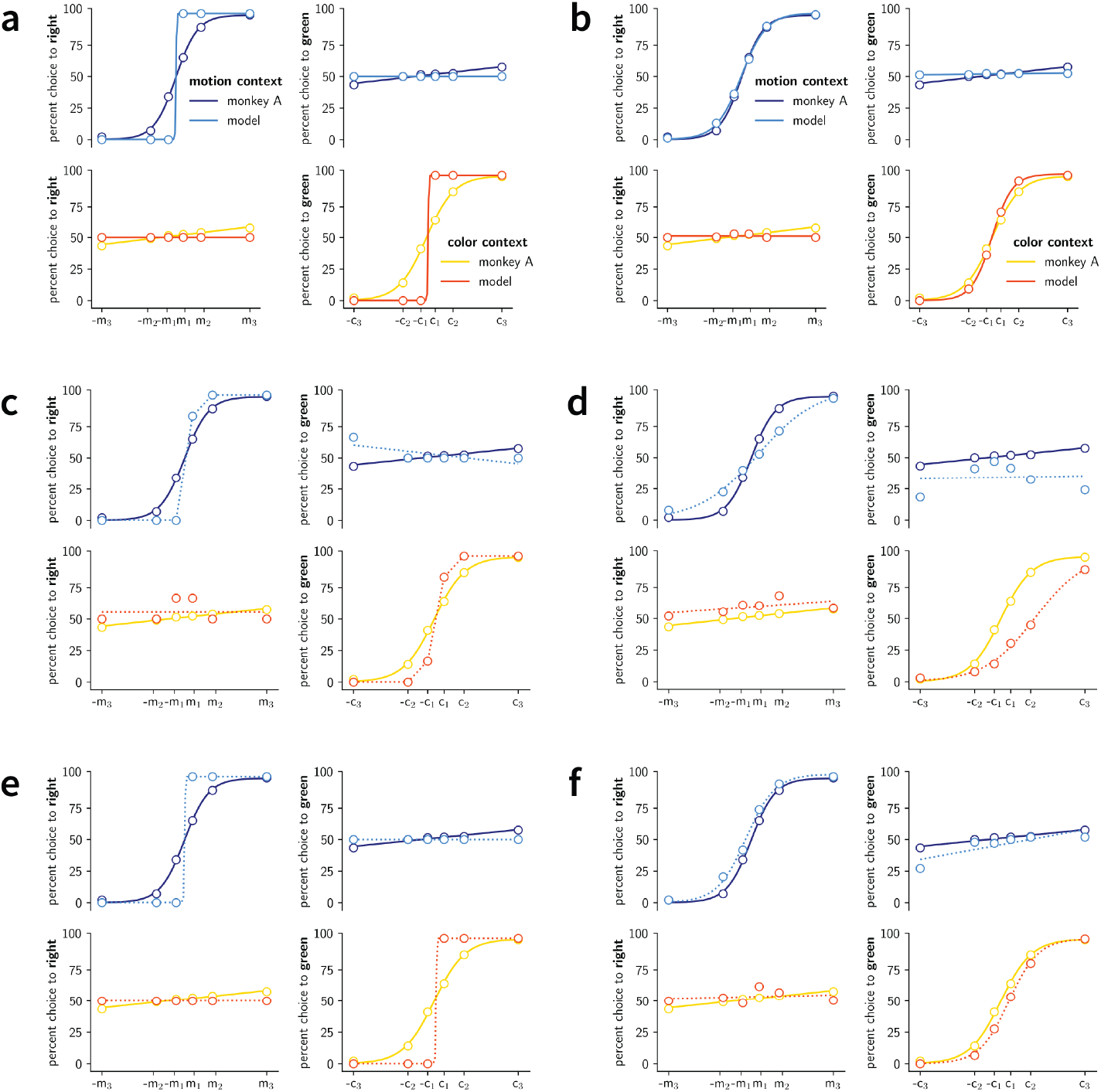
Psychophysical response of candidate models with and without noise corruption of the stimulus. **a.** The model that is the main focus of this paper is able to perform perfect discrimination on trials of all coherence values, which is inconsistent with the behavioral response we see in live subjects engaged in the same perceptual discrimination task. The model also demonstrates perfect “indifference” to the coherence values of the irrelevant stimulus in any given context. This, too, is inconsistent with biological behavior, which shows evidence of extremely minor—but nonetheless existent—preference for stronger coherence values of the irrelevant stimulus. **b.** Figure from the main text. **c.** The psychophysical response of the candidate comparison model with no hidden neurons when the stimulus is not corrupted with noise, and **d.** is the psychophysical response of the same model when the stimulus has the same level of corruption as in **b. e.** The psychophysical response of the candidate comparison model with hidden neurons but in which the input stimulus is projected to the entire population when the stimulus is not corrupted with noise, and **e.** is the psychophysical response of the same model when the stimulus has the same level of corruption as in **b, d.** The psychophysical performance for the second candidate model closely approximates the behavior of the monkey, but because the PSTHs of the observed population bear little likeness to the target experimental dataset, we opted not to use this model.

## B.3 State space dynamics of model trained only to solve the behavioral task

**Figure 9:**
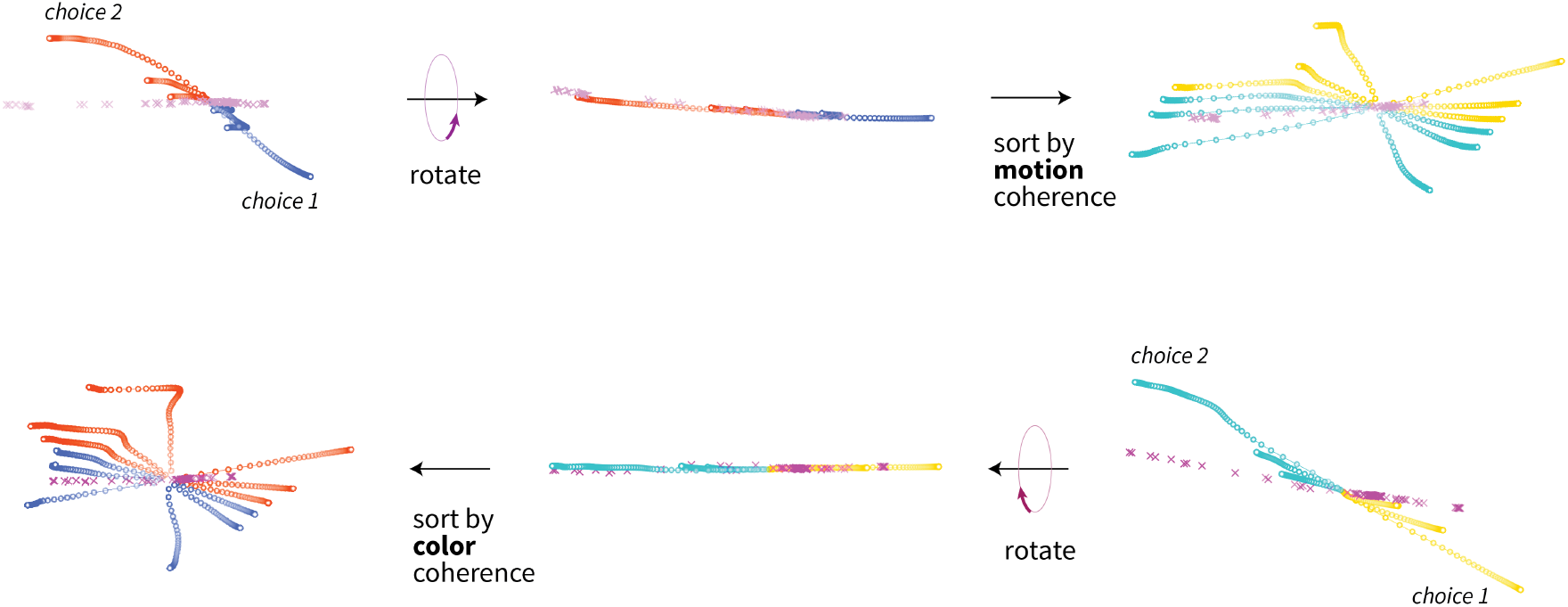
State space dynamics of model trained only to solve the behavioral task. Using the model described in Appendix A, we trained a network to solve the same discrimination task that our model network is trained to solve, except that we did not require the network to reproduce experimental data, as well. We did this in order to confirm that the low-dimensional schematic governing input integration in our model network—particularly its difference to that propose in [4]— was not simply a result of using a different training method (back propagation through time, as in [4], versus our adapted FORCE/RLS rule). The dynamics of this model are not *identical* to those of the model used in [4], although they do follow a similar low-dimensional process, governed by a context sensitive line attractor and associated selection vectors. The dynamics shown here are the result of training a network with 800 neurons. The top row displays dynamics in the color context along the line attractor spanned by slow points in the state space, rotated, and then sorted by the coherence of the irrelevant stimulus value. The bottom row shows the same thing, but in the motion context. Colors used here correspond to those used in *Figure 2*, with corresponding behavioral choices labeled. The axes of the state space are as defined in Chapter 4 (using the right-zero eigenvector to define the choice axis, and using the color and motion filter weights of the input matrix to the network as the color and motion axes, respectively).

## c Methods

## C.1 Primary network architecture

For all experiments, we start with a randomly initialized, strongly recurrent neural network (schematic in *Figure 1*). The activities of neurons in the network are described by:

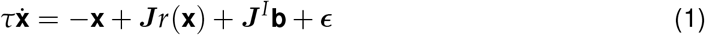

where *τ* is a time constant, *x*_*i*_ is the activity of neuron *i*, ***J*** is a recurrent weight matrix such that *J*_*ij*_ describes the strength of connection between neurons *i* and *j*, neuron *i* is presynaptic to neuron *j*, and 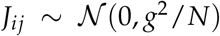, where *g* is a hyperparamter, the magnitude of which determines the strength of connections between neurons in the network, and *N* is the number of neurons in the network. At each time step, the activity of the network from the previous time step is introduced to the network via a nonlinear function *r*(·). In order to restrict neural activity between a plausible firing rate range of *r*(·) ∈ [0, 1], we use the non-linear activation function *r*(**x**) = 1/1 + *e*^−*x*^ applied element-wise to **x**. The magnitude of the stimuli introduced to neuron *i* is determined by (***J***^*I*^**b**)_*i*_, and 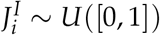. Finally, ***ϵ*** is a vector of stochastic noise introduced directly onto each neuron, and 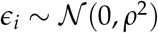. For most experiments, we used *ρ* ∈ [0, 1].

The model population can be partitioned into two subsets: hidden and observed neurons. Observed neurons are trained (the procedure is described in detail below) to represent a target function, while hidden neurons have no associated target function. Inputs to the network are restricted to a sparse subset of the hidden population. If there are *N* observed neurons, *M* hidden neurons, and *P* inputs to the network, we can think of ***J***^*I*^ as the concatenation of two matrices: an *N* × *P* matrix of all zeros, and an *M* × *P* matrix, a fraction of whose entries are 0; the rest of which are as defined above. In this structure, inputs largely drive the hidden population, which then in turn drives the observed population via a transformation of the input signal. As stated in the text, the hidden population maintains a feed-forward connection into the observed population, is not itself recurrently connected, and has no associated target function.

For each network simulation, we solved the above network equation over some time interval *δt* using Euler integration:

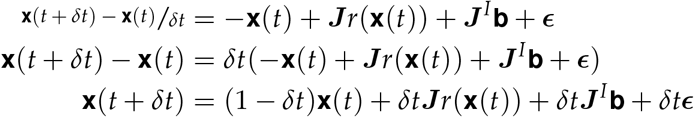

for time *t* ∈ [0, *T*) for some predetermined time interval size *T*. The readout at each time step is simply *r*(**x**(*t*)). In other words, the readout of the network at any time step is the firing rate of each neuron. This allows the observed state of the network to be the direct firing patterns of each neuron in the network, rather than a linear transformation of the network activities over an external set of learned weights.

For modeling the output of both neural data and the target behavioral task variable, we define a final node in the network, which we denoted as *z*, that is virtually external, but still within the network. This node is defined as

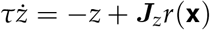

and is trivially solved, like above, as *z*(*t* + *δt*) = (1 − *δt*)*z*(*t*) + *δt****J***_*z*_*r*(**x**(*t*)). Here, ***J***_*z*_ is the set of weights representing the presynaptic strength of each neuron in the network onto the readout neuron, indexed with *z*. Effectively, the behavioral node’s readout is the weighted sum of the non-linear activations of the nodes in the network at each time step, but its dynamics are not explicitly reintroduced into the other nodes in the network at each time step. In other words, the behavioral node is “aware” of other nodes’ readouts in the network, but other nodes are unaware of its state.

In order to model context-dependent feature integration task outlined in [4], we used stimulus vectors **b** = (cc mc ac am ret)^T^, where cc, mc ∈ {−0.5, −0.17, −0.05, 0.05, 0.17, 0.5}, and ac = 1, am = 0 in the color context, and ac = 0, am = 1 in the motion context. ret is set to zero during stimulus presentation, and is set to 1 to indicate to the network when a trial has ended. The target behavior includes reporting the sign of the relevant stimulus. For example, if am = 1, and mc = 0.5, we would expect the network to report that it has integrated “positive” motion coherence, by reporting *z* = 1.

In some trials, we corrupt the input stimulus with stochastic noise in order to simulate the noisy nature of both the stimuli in the experimental setting and corruption of stimuli hitting the retina of an experimental subject (due to a variety of factors including the refraction of light by retinal fluids, etc.). For trials in which the input stimulus is corrupted by stochastic noise, the stimulus values become time-varying functions:

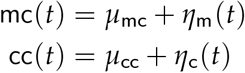

where *μ*_mc_, *μ*_mc_ ∈ {−0.5, −0.17, −0.05, 0.05, 0.17, 0.5} and 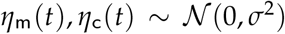. Then, 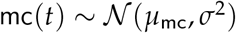 and 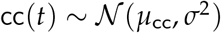. The input vector then becomes **b** = (cc(*t*) mc(*t*) ac am ret)^T^. For experiments, we typically restricted the variance of the stochastic noise to *σ*^2^ ∈ [0, 1.5].

We use this model structure for several reasons. Principally, this model has very few “built-in” features, which allows it to act as a general model of population-level neural computation. The network is not designed for any one specific task, and thus, like the brain, is able to perform a wide range of tasks that are mutually unrelated. Ultimately, we are looking to deduce the underlying structure of how context-dependent information is represented in the brain: the recurrent weights of the RNN simulate recurrent connectivity between neurons in the biological brain, allowing neurons in one area of the network to communicate with other ones, particularly over time.

## C.2 Training procedure

The FORCE [28] training technique attempts to minimize a weighted linear least squares cost function with respect to some target function *f* (*t*). Here, *f* (*t*) is a column vector in which the *i*-th entry describes the firing rate of neuron *i* in the population of recorded neural data at some time *t*. For our purposes, we define the cost function as:

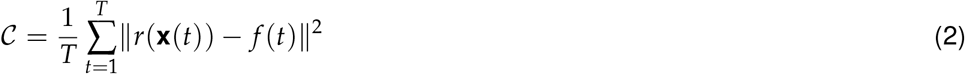

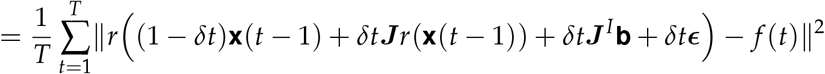

where || · || is the 2-norm. The task, then, is to train individual neurons in the network to match their activities such that the firing rates are as close as possible to the target firing rates from the recorded neural activities. The FORCE algorithm accomplishes this by learning the optimal weights of ***J***, the recurrent connectivity matrix, using the recursive least-squares filter (RLS). RLS, however, would appear unfit for the task of learning weights to minimize 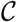, as RLS is a training procedure for learning weights to minimize a linear least squares objective function (the network readout, by definition, is a non-linear function of the neuron activities). The use of FORCE/RLS is preferred, mainly for computational efficiency, and so we rewrite the cost function in equivalent linear terms:

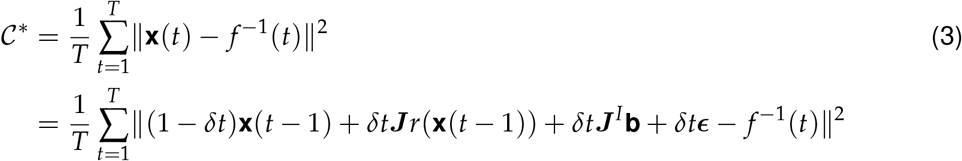

where *f*^−1^(*t*) := *r*^−1^(*f* (*t*)), or the target function passed through the inverse of the non-linear activation function applied to the neurons at each readout step. In this case, *r*^−1^(*x*) = ln(*x*/1 − *x*), which is applied element-wise to the target vector. Observe that this objective is certainly a weighted linear least squares function; the only other difference, then, between 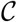 and 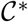 is that 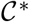 describes a loss between the activities of each neuron (rather than the firing rates) and the target function passed through the inverse of the activation function (from which we surmise the firing rates of neurons in the network) rather than between the firing rates of each neuron and the corresponding target function. If we compose both components of the cost function (**x**(*t*) and *f*^−1^(*t*)) with the non-linear activation function, *r*(· ), we recover the original cost function. The filter seeks to elect weights for ***J*** that minimize 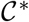 (and, hence, also 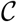), which is done by first calculating the error signal ***e***(*t*) between the transformed target function *f* −1(*t*) and activities each neuron in the network **x**(*t*),

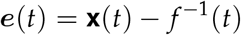

updating the cross-correlation matrix of the firing rates, ***P***(*t*),

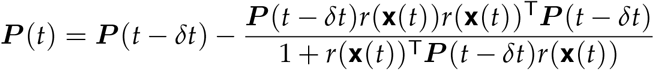

and finally adjusting weights of the recurrent connectivity matrix ***J***:

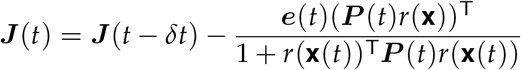

For these experiments 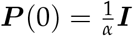, where *α* is considered the approximate learning rate of the FORCE procedure (we typically used an *α ≪ N*, the number of neurons in the network, usually using *α* = 1). For each update to the recurrent weight matrix, we restricted modifications to connections presynaptic to the pinned neurons in the network, or the neurons which had a direct target neuron on which its output was trained. The weights presynaptic to hidden neurons in the population whose firing rates were not recorded were not modified.

The error signal, which is reintroduced into the network via the recurrent weight update, is kept relatively small throughout the training procedure (the mathematical basis for this phenomenon is described in [28]). This fact is important for two reasons: (1) such an error control allows for fast convergence in the task of learning optimal recurrent weights ***J***, which is important when approximating high-dimensional target functions that carry with them high computational costs in modeling, and (2) because the error signal re-introduced to the network does not induce chaotic behavior in the RNN, which is often a challenge one must surmount in training RNNs to perform complex, temporally dependent tasks.

## C.3 Task-based network architecture

The model structure is similar to that used in this paper. The activities of neurons in the network are described by:

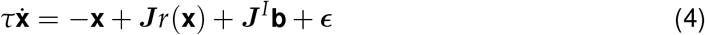

where *τ* is a time constant, *x*_*i*_ is the activity of neuron *i*, ***J*** is a recurrent weight matrix such that *J_ij_* describes the strength of connection between neurons *i* and *j*, neuron *i* is presynaptic to neuron *j*, and 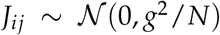, where *g* is a hyperparamter, the magnitude of which determines the strength of connections between neurons in the network, and *N* is the number of neurons in the network. At each time step, the activity of the network from the previous time step is introduced to the network via a nonlinear function *r*( · ). We require no restriction on the firing rate range of a neuron ( as all neurons are effectively hidden), and so we use the non-linear activation function *r*(**x**) = tanh(**x**) applied element-wise to **x**. The magnitude of the stimuli introduced to neuron *i* is determined by (***J***^*I*^**b**)_*i*_, and 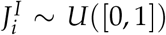. Finally, ***ϵ*** is a vector of stochastic noise introduced directly onto each neuron, and 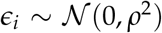. For most experiments, we used *ρ*^2^ ∈ [0, 1].

Unlike the model that is the focus of this paper, there are two sets of trainable weights. One set defines the recurrent connections within network the network, and the other linearly transforms the firing rates of neurons in the network to produce a target variable:

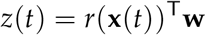

Similar to the network that is the focus of this paper, in order to model context-dependent feature integration task outlined in [4], we used stimulus vectors **b** = (cc mc ac am ret)^T^, where cc, mc ∈ {−0.5, −0.17, −0.05, 0.05, 0.17, 0.5}, and ac = 1, am = 0 in the color context, and ac = 0, am = 1 in the motion context. ret is set to zero during stimulus presentation, and is set to 1 to indicate to the network when a trial has ended. The target behavior includes reporting the sign of the relevant stimulus. For example, if am = 1, and mc = 0.5, we would expect the network to report that it has integrated “positive” motion coherence, by reporting *z* = 1.

## C.4 Task based training method

Here, too, we employ FORCE learning [28], though 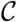 takes a different form:

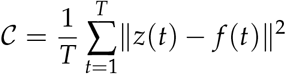

Here, we learn weights on **w** using the RLS filter. The filter seeks to elect weights **w** that minimize 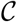, which is done by first calculating the error signal *e*(*t*) between desired output *f*(*t*) and network output *r*(**x**(*t*))**w**(*t* − *δt*),

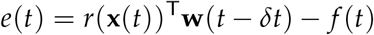

and using the calculated error signal *e*(*t*) to update the weights at each iteration **w**(*t*),

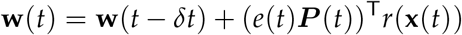

and then finally, updating the cross-correlation matrix ***P***(*t*),

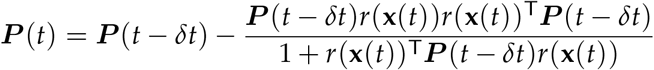

For the experiments detailed below, we employed a variant of the FORCE learning scheme that also adjusts weights of the connectivity matrix ***J***:

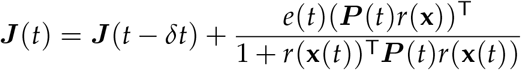

For these experiments, 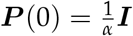, where *α* is considered the approximate learning rate of the FORCE procedure (we typically used an *α ≪ N*, the number of neurons in the network, usually using *α* = 1). Additionally, **w**(0) = **0**.

## C.5 State space analysis, or reverse-engineering the network

## C.5.1 Theoretical background and methods

A significant portion of the analysis of our model involves treating the RNN as a globally non-linear dynamical system approximated by a collection of local linear systems surrounding fixed and semi-stationary points in the state space of the RNN. By studying the system in the locally linear regimes, which are analyzable in ways that the global system is not, we gain intuition for how the network attempts to solve the larger goal of integrating information and dynamically representing that information based on a context.

## C.5.2 Locating fixed points in the model state space

The process of finding a fixed point in the system amounts to identifying a location in the state space for which small perturbations along any of the spanned axes results in minimal movement of the system from the point. The dynamical system is globally described by *Equation 3.1*, which we rewrite in compact notion: 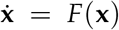. We want to find a set of points *X*^*^ = {**x**^*^ | *F*(**x**^*^) = **0** ∨ *F*(**x**^*^) ≈ *F*(**x**^*^ + *δ***x**) for some sufficiently small *δ***x**. The former condition, *F*(**x**^*^) = **0**, describes a strict fixed point. The complexity of the dynamical system modeled by the RNN requires us to also consider slow points where dynamics are not necessarily *fixed*, but which nonetheless behave like strictly fixed points within bounded regions (captured by the latter condition, *F*(**x**^*^) ≈ *F*(**x**^*^ + *δ***x**)).

To find these points, we define an auxiliary function: 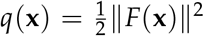, which can be thought of as, effectively, the kinetic energy of the system *F*(·) at some point **x**. We use this function because it is a scalar function, and thus is amenable to optimization software. Because *q*(·) is a sum of squares, we know that it reaches its minimum wherever *F*(·) does, and so by finding where *q*(·) reaches 0, we find where *F*(·) = **0**.

## C.5.3 Locally linear dynamics

Consider the Taylor expansion of 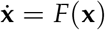 around some fixed point **x**:

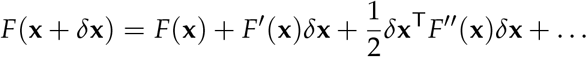

Because the point **x** is a fixed point, *F*(**x**) = **0**, and the second order term of the Taylor series (and all terms succeeding it) are approximately **0**, so the magnitude of first order term dominates all other terms, or else 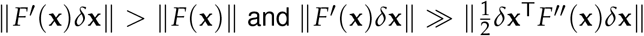 (we are concerned primarily with perturbations *δ***x** that are quite small, which substantially reduce the magnitude of the second-order term). These inequalities illustrate why some points **x** such that *F*(**x**) = **0** are ideal candidates for linearization: they afford us a lower bound on ||*δ***x**||, which is 0. In other words, all infinitesimally small perturbations ||*δ***x**|| > 0 around some points **x** such that *F*(**x**) = **0** will generate local linear or approximately linear dynamics.

Because we examine dynamics around fixed points, or points where *F*(**x**) = **0**, our expansion of *F*(**x**+ *δ***x**) reduces to:

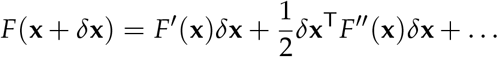

Let ***p*** = *δ***x**. Heeding the second inequality 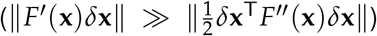, we omit the second order term of this expansion, which yields:

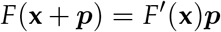

which is the general form for a linear dynamical system: 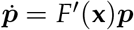, which is linear in ***p***. Thus, when studying local linear dynamics around some fixed point **x**, our analysis will primarily be concerned with the matrix ***M*** = *F*′ (**x**), the Jacobian of *F*(·). In fact, we can think of ***M*** as the linear approximation of the network around fixed points, which aids in the following analysis.

We are interested in understanding local linear dynamics around ***M***, and so we perform an eigende composition on ***M*** :

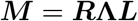

such that ***L*** = ***R***^−1^, and where the *i*-th row of ***L*** is the *i*-th left eigenvector of ***M*** , the *i*-th column of ***R*** is the *i*-th right eigenvector, and **Λ** is a diagonal matrix such that **Λ**_*ii*_ is the *i*-th eigenvalue corresponding to the *i*-th eigenvectors of ***L*** and ***R***. Let *λ*_*i*_ := **Λ**_*ii*_, and let **l***i* and **r***i* denote the *i*-th left and right eigenvector, respectively. Then, **l**_*i*_***M*** = *λ*_*i*_**l**_*i*_ and ***M*r**_*i*_ = *λ*_*i*_**r**_*i*_. Such a decomposition offers tools to study the evolution of the system according to independent bases, which drastically simplifies the analysis of the RNN around a fixed point.

Then, returning to the equation describing the locally linear dynamical system:

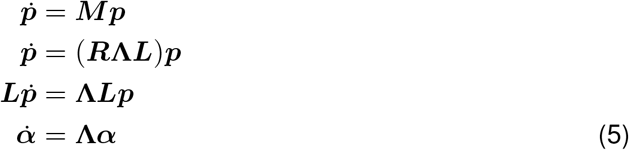

where ***α*** := ***Lp***. Then any mode *i* can be described by 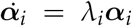 in this locally linear regime, and, because the system is described over the diagonalized left eigenbasis, mode *i*’s evolution in the regime is independent of all other modes’ states in the system. Isolating the activities of each mode independently affords us a means of characterizing high-dimensional systems based on the activities of a subset of modes within the system, particularly when the contribution of this subset of modes to the evolution of the system is significantly larger than others.

*Equation 4.1* describes the dynamics of a general, locally linear dynamical system around a stationary point. Such an analysis is still applicable to our system of interest (an RNN responding to incoming sensory evidence), albeit with an extension. In describing our system’s evolution through the repeated application of the left eigenvectors associated with the Jacobian of the network around a fixed point, we can argue that the incorporation of sensory evidence into the network amounts to adding to the network state the projection of the stimulus of interest, filtered through the input weights associated with that stimulus, onto the left eigenbasis. Additionally, because each mode of *Equation 4.1* evolves independently, we can simply add this projection to each mode.

In the case supposed in [4], fixed points and the locally linear evolutions within the state space around these points organize in a line attractor, or a line in the state space along which fixed and slow points lie. For this to occur, the spectra of ***M*** at all fixed or slow points must have a specific structure: max(**Λ**) ≈ 0 and the other *N* −1 eigenvalues should have very large negative real parts (or, indexing eigenvalues by their magnitude, |*σ*_*i*_| ≫ |*ω*_*i*_| for *i* = 2, … , *N*)^2^.

Along the *N* −1 modes with eigenvalues that have large negative real parts, trajectories rapidly decay; only along the mode with the near 0 eigenvalue does the system integrate over the course of the trajectory. Let **l**_0_, **r**_0_ denote the left and right eigenvectors of the 0 mode. The system evolution along this mode is observable as the result of a projection of the trajectory along **l**_0_ onto the **r**_0_. Then, the line attractor is spanned **r**_0_.

## C.6 Transforming neurological data into a target function

Our target neural data set comes from Mante et al.’s study, and is comprised of neural recordings taken from FEF of two rhesus monkeys trained to perform the context-dependent feature discrimination task described in Chapter 2. We followed Mante et al.’s procedure for parsing, cleaning and organizing the data, which we describe briefly in this section.

For their experiments, Mante et al. taught the task to two rhesus monkeys, labeled A and F, and performed several, randomized trials in each recording session with both A and F. Each recording session is contained within its own .mat file; each one contains, on average, 1,200 trials for each unit. Each trial is comprised of a neural recording, lasting 850 ms, and values indicating the motion coherence of the dots on the display, the color coherence of the dots, the context of the trial (what feature the monkey was instructed to attend to), and whether or not the monkey correctly completed the trial by directing a saccade to the target.

The neural recordings from the experiments take the form of a sequence of 0’s and 1’s, one in each time bin. In each bin, 1 indicates detected electrophysiological activity, and 0 indicates the absence of any detected activity. Mante et al. performed their recordings with the hopes of analyzing neural behavior at the neuron population level; while this method allows for the analysis of neural dynamics across larger neural landscapes and greater insight into how populations of neurons act during activities of interest, this method of recording is also far less precise than single-neuron recordings. This results in remarkably high variability in the recorded neural response across trials in a single unit for which the monkey was presented the exact same information. This fact, along with the underlying assumption in our RNN model that functions it is trained to reproduce are continuous (unlike the binary nature of the neural recordings) requires us to manipulate the neural data in a way that preserves its structure but that also makes it more amenable to reproduction by our model.

For each unit, we averaged the neural recordings across conditions. We identified 144 possible conditions for the data: some permutation of 6 possible motion coherence values and 6 possible color coherence values, 2 possible contexts (attend to color coherence or attend to motion coherence) and 2 possible outcomes of the experiment (either the monkey got it right or wrong). The experimental data for each unit had roughly the same number of trials for each of these conditions, which made averaging across them a reasonable estimate for neural activity in that unit in any particular condition. For each time bin in each condition, we calculated the average strength of the neural pulse by summing the neural activations across each time bin for each condition and divided by the number of trials in each respective condition. We recovered the firing rate of individual neurons by using a moving box average with a width of 50 bins. We focused only on neural traces 100 ms after stimulus onset, leaving a trace 750 ms in length.

The moving box average resulted in traces that were 750 / 50 = 15 bins long.

The final step involved smoothing and re-sampling the box-averaged neural response. We first smoothed the response using a Gaussian kernel (*σ* = 40 ms) and then re-sampled this smoothed signal to create a sequence of 200 bins. After smoothing and re-sampling, we re-shaped the data, by z-scoring the value in each time bin and throughout each condition by subtracting the mean of the total average activity over the entire unit, and dividing by the standard deviation of the activity over the entire unit.

## D Statistics

We use two primary statistics to quantify the accuracy of our model in simulating neural trajectories: 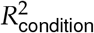 and 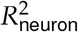.

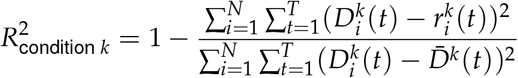

where *N* is the number of observed neurons in the system, *D*_*i*_(*t*) is the target value of neuron *i* at time *t* in condition *k*, where a condition is a set of motion and color coherence values, and an associated context cue, 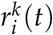 is model neuron *i*’s output at time *t* in condition *k*, and 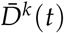 is the mean firing rate across all neurons in condition *k* at time *t*. 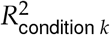 is the *R*^2^ coefficient, specifically, for describing the correlation between the target function and the output of the model for some *condition k*.

We are also interested in quantifying neuron-to-neuron likeness across conditioned trajectories, and so we also use 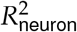 :

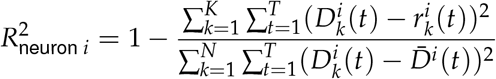

where *K* is the total number of conditions, 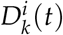 is the target value of neuron *i* in condition *k* at time *t*, 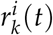 is model neuron *i*’s output at time *t* in condition *k* and 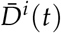 is the mean firing rate of neuron *i* across all conditions at time *t*.

## Footnotes

1 Because the weights within the hidden population are not modified during training, we omit these in this figure.

2 An important note here is that the dimensionality of the attractor is intimately linked to the number of near-zero eigenvalues in the spectrum of ***M***. If the system organized points along a plane attractor and dynamics along that plane were stable or near-stable, the spectrum of ***M*** would contain 2 near-zero eigenvalues, and so on for higher-dimensional attractor manifolds.

